# Homeobox transcription factor MSX2 down-regulates ERα action to facilitate anti-estrogensensitivity in ERα+ breast cancer

**DOI:** 10.64898/2026.06.12.731842

**Authors:** Jianwei Feng, Kai Zeng, Anqi Wang, Hongmiao Sun, Zining Jin, Zhan Liu, Shu Feng, Xiaodi Sun, Yu Bai, Mingcong He, Wensu Liu, Manlin Wang, Dongjun Yang, Jin Wang, Wei Liu, Lu Xu, Chunyu Wang, Shengli Wang, Yue Zhao

## Abstract

Breast cancer is the most common malignancy among women worldwide. Estrogen receptor α (ERα)-positive breast cancer is about 70% in the total breast cancer. It has variable prognosis and high recurrent risk, frequently developing resistance to endocrine therapies. Herein, we have identified that a homeobox transcription factor MSX2 downregulated in ERα-positive breast cancer is correlated with the advanced grade and poor survival outcomes. Our results have demonstrated that MSX2 interacts with ERα to inhibit ERα-mediated transactivation. Mechanistically, MSX2 associates with HDAC1/HDAC2 to form a protein complex. MSX2 is required for recruitment of HDAC1/HDAC2 complex to estrogen response elements (EREs) region on ERα downstream target genes, reducing histone H3K9ac and H3K27ac levels to inactivate gene transcription. Functionally, ectopic expression of MSX2 suppresses cell growth and enhances the sensitivity to anti-estrogen treatment in ERα-positive breast cancer cells. Our results indicate that MSX2 as a novel co-repressor of ERα is involved in suppression of ERα-positive breast cancer progression. Restoring MSX2 protein may provide a promising strategy to overcome endocrine resistance in ERα-positive breast cancer.

## Introduction

Breast cancer (BCa) remains the most frequently diagnosed malignancy and a leading cause of cancer-related mortality among women worldwide, with an estimated 2.3 million new cases and 670,000 deaths reported in 2022(Bray *et al*, 2024). Based on immunohistochemical (IHC) profiles of estrogen receptor (ER), progesterone receptor (PR), HER2, and Ki-67, breast cancer is classified into several molecular subtypes, including Luminal A, Luminal B, HER2-positive, and triple-negative breast cancer (TNBC)(Goldhirsch *et al*, 2013). The Luminal subtype, the most common breast cancer type characterized by the expression of estrogen receptor alpha (ERα), has variable prognosis and high recurrent risk(Ignatiadis & Sotiriou, 2013; Siegel *et al*, 2024). As a ligand-dependent transcription factor, ERα drives the transcription of target genes in response to estrogen (E2), thereby promoting the proliferation, differentiation, and metastasis of mammary epithelial cells(Clusan *et al*, 2023; Green & Carroll, 2007; Ozyurt & Ozpolat, 2022; Silva-Cázares *et al*, 2024).

Given its central role in BCa progression, ERα is the primary target of endocrine therapies, including selective estrogen receptor modulators (SERMs, e.g., tamoxifen), selective estrogen receptor degraders (SERDs, e.g., fulvestrant), and aromatase inhibitors (AIs)(Kim & Lukong, 2025; Patel & Bihani, 2018). Endocrine therapy combined with CDK4/6 inhibitors is standard treatment for luminal breast cancer (Chong *et al*, 2020; Morrison *et al*, 2024). However, the development of primary and acquired resistance to these therapies poses a major clinical challenge. The mechanisms underlying resistance are complex and multifaceted, involving *ESR1* mutations, loss of ERα expression, aberrant crosstalk between ERα and multiple signaling pathways (e.g., growth factor pathways), dysregulation of ERα transcriptional control (including altered coregulator activity), and dysregulation of cell cycle checkpoints(Altwegg & Vadlamudi, 2021; Bower *et al*, 2017; Clusan *et al*., 2023; Garcia-Martinez *et al*, 2021; Jeselsohn *et al*, 2015; Liang *et al*, 2025; Ma *et al*, 2022; Yan *et al*, 2024). Therefore, understanding ERα transcriptional regulation is critical for developing strategies to overcome endocrine resistance.

ERα function is tightly controlled by co-regulators that assemble into multi-protein complexes to activate or repress transcription of target genes (Altwegg & Vadlamudi, 2021). Identified coactivators include p160 family (SRC-1, SRC-2, SRC-3), SWI/SNF chromatin remodeling complex, p300/CBP, and histone methyltransferases(An *et al*, 2004; Johnson & O’Malley, 2012; Luan *et al*, 2024; Sun *et al*, 2020; Yi *et al*, 2015), Conversely, corepressors, such as BRCA1, SMRT, NCOR1, LCoR, and YAP et al., inhibit ERα activity or promote its degradation(Altwegg & Vadlamudi, 2021; Britschgi *et al*, 2017; Dobrzycka *et al*, 2003; Fan *et al*, 1999). Dysregulation of the co-regulators disrupts ERα-driven transcriptional programs, contributing to tumor progression and therapy resistance. Thus, identification of the novel co-regulators of ERα and understanding well the multifaceted modulation of ERα signaling pathways would provide the new therapeutic strategies for the advanced luminal breast cancer.

MSX2, a homeobox transcription factor located at 5q34–q35, is a key regulator of morphogenesis in diverse tissues, including craniofacial bones, mammary glands, and hair follicles (Bendall & Abate-Shen, 2000; Jabs *et al*, 1993; Ma *et al*, 2003; Yu *et al*, 2019b). Beyond morphogenesis, MSX2 is critical for embryonic and extraembryonic development, as well as stem cell fate determination. For instance, in placental development, the SP6–MSX2 axis, activated by SP6 and p300 at H3K27ac-marked enhancers, is essential for maintaining trophoblast stem cell identity by preventing premature differentiation(Chen *et al*, 2024). Mechanistically, MSX2 partners with the canonical BAF (cBAF) subcomplex of the SWI/SNF chromatin remodeling complex to repress differentiation-associated genes, with its loss leading to increased chromatin accessibility and enhancer activation(Hornbachner *et al*, 2021). In pluripotent stem cells, MSX2 drives mesendodermal lineage specification by repressing SOX2 and activating NODAL, forming a mutually antagonistic loop with SOX2 that fine-tunes cell fate(Wu *et al*, 2015).

With its established functions in tumor development and stem cell programs, MSX2 has been recognized as a context-dependent regulator in cancer. It functions as either an oncogene or a tumor suppressor in different tissue-derived cancers. In ovarian endometrioid adenocarcinoma, pancreatic cancer, and colorectal cancer, MSX2 exerts oncogenic functions by promoting epithelial-mesenchymal transition (EMT) and chemoresistance(Liu *et al*, 2017; Satoh *et al*, 2008; Zhai *et al*, 2011). In ovarian endometrioid adenocarcinoma, activating downstream of the Wnt/β-catenin-TCF signaling pathway, MSX2 promotes epithelial-mesenchymal transition (EMT), enhancing tumor invasion and metastasis(Zhai *et al*., 2011). In pancreatic cancer, it significantly promotes chemoresistance by upregulating drug resistance genes, such as MRP2 and ABCG2(Hamada *et al*, 2012). Conversely, MSX2 displays tumor-suppressive properties in osteosarcoma and melanoma(Gremel *et al*, 2011; Wu *et al*, 2021). In osteosarcoma, MSX2 directly binds to the SOX2 promoter and represses its transcription, thereby diminishing tumor cell stemness, proliferation, metastatic potential, and increasing chemosensitivity(Wu *et al*., 2021). And in melanoma, it downregulates anti-apoptotic proteins Bcl-2 and Survivin, upregulates the cell cycle inhibitor P21, and suppresses tumor cell proliferation, invasion, migration, stemness, while promoting apoptosis(Gremel *et al*., 2011). MSX2 exhibits a dual role in cancer, driving malignancy in some contexts (e.g., ovarian, pancreatic cancers), while suppressing the tumor progress in others (e.g., osteosarcoma, melanoma). The previous study has shown that ectopic expression of MSX2 inhibits cell proliferation and leads to induction of apoptosis in breast cancer cells (Lanigan *et al*, 2010). However, what is molecular mechanism underlying the biological functions of MSX2 on ERα-positive breast cancer progression is still elusive.

In this study, we have identified that MSX2 is significantly down-regulated in clinical breast cancer specimens and its lowly expression is closely associated with patient endocrine therapy resistance status. Furthermore, MSX2 physically interacts with ERα in breast cancer cells, significantly inhibiting ERα-induced transcriptional activation. MSX2 is recruited to the promoter regions of ERα downstream genes. Strikingly, MSX2 facilitates the recruitment of HDAC1/HDAC2-containing protein complex at estrogen response elements (EREs) to reduce H3K27ac and H3K9ac levels, maintaining an inactive chromatin state. Importantly, ectopic expression of MSX2 significantly inhibits cell proliferation and synergizes with anti-estrogen to enhance endocrine treatment sensitivity in breast cancer. Collectively, our findings indicate that MSX2 acts as a critical epigenetic repressor of E2/ERα oncogenic signaling pathway, proposing that upregulation of MSX2 expression may provide a novel strategy to overcome endocrine resistance in ERα-positive breast cancer.

## Results

### MSX2 is identified as an endocrine response–associated regulator of tumor progression and proliferative signaling in breast cancer

To identify novel factors potentially associated with endocrine resistance in ER-positive breast cancer, we performed an integrated transcriptomic analysis of multiple independent datasets representing tumorigenesis and acquired resistance. Tamoxifen resistance–related genes (GSE26459), fulvestrant resistance–related genes (GSE190384), and ER-associated genes in the TCGA -BRCA cohort were intersected. This analysis yielded 34 overlapping candidates, including MSX2 and several established luminal/ER-related genes, supporting the relevance of MSX2 to endocrine response and ER-associated biology (Figure 1A). In addition, we examined the patient cohort dataset GSE9195, which consisted of tamoxifen-treated ER-positive breast cancer patients, and found that MSX2 was downregulated in resistant samples compared with sensitive samples, suggesting a potential association between MSX2 loss and endocrine resistance (Figure 1B). To assess the clinical relevance of MSX2, Kaplan–Meier survival analysis demonstrated that patients with low MSX2 expression exhibited significantly poorer survival compared with those with high expression (Figure 1C). Consistently, analysis of the METABRIC cohort revealed that MSX2 expression progressively decreased with increasing tumor grade, indicating its association with tumor progression (Figure 1D). To further explore the biological processes associated with MSX2, gene set enrichment analysis (GSEA) was performed. Tumors with low MSX2 expression showed significant enrichment of E2F1 target genes and G2M checkpoint pathways, indicating enhanced cell cycle activity upon MSX2 downregulation (Figure 1E–F). In addition, correlation analysis revealed that MSX2 expression was inversely associated with MYC and E2F1, a representative ERα downstream target gene involved in cell proliferation, suggesting that reduced MSX2 expression may be linked to increased ER-driven transcriptional output (Figure 1G). Collectively, these results identify MSX2 as an endocrine response–associated candidate in breast cancer and suggest that its downregulation is associated with poor prognosis, tumor progression, and activation of proliferation-related transcriptional programs.

**Figure 1.**
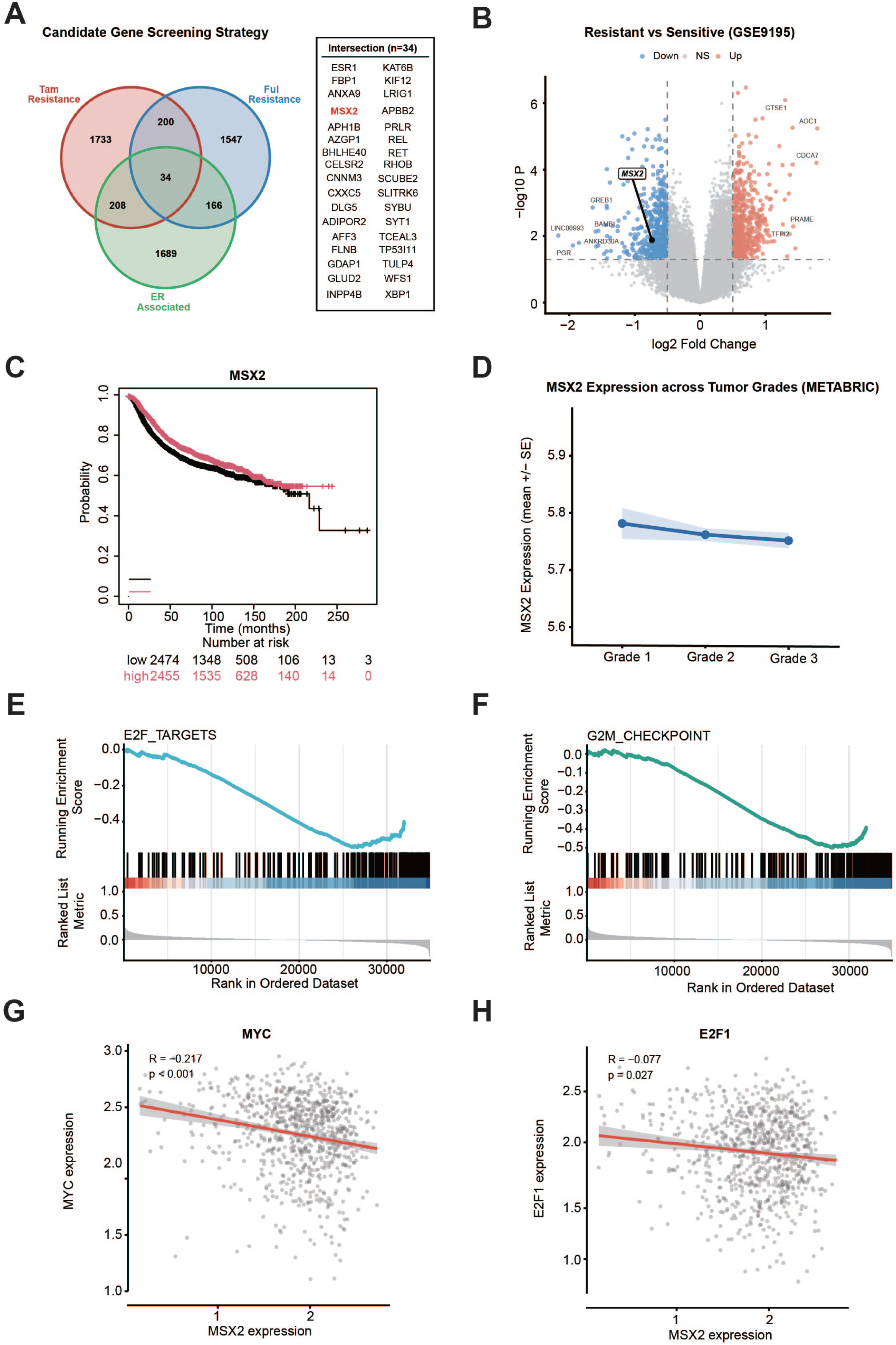
Bioinformatic identification of MSX2 as an endocrine response–associated regulator in breast cancer. **A**: Venn diagram showing the overlap among tamoxifen resistance–related genes, fulvestrant resistance–related genes, and ER-associated genes, identifying 34 candidate genes including MSX2. **B**: Volcano plot showing that MSX2 is downregulated in endocrine-resistant samples compared with sensitive controls. **C**: Kaplan–Meier survival analysis showing that low MSX2 expression is associated with poor prognosis in breast cancer patients. **D**: MSX2 expression across tumor grades, showing a progressive decrease with increasing tumor grade. **E–F**: Gene set enrichment analysis demonstrating enrichment of cell cycle–related pathways, including E2F targets and G2M checkpoint, in tumors with low MSX2 expression. **G**: Correlation analysis showing a significant inverse association between MSX2, MYC and E2F1 expression. Samples were stratified into MSX2 high and low expression groups based on the median expression value.

### Downregulation of MSX2 is correlated with the poor prognosis in ERα+ breast cancer

The above data analysis indicates that MSX2 is associated with resistance to endocrine therapy in breast cancer, and that high MSX2 expression correlates with a favorable prognosis. Next, we aim to further verify the expression of MSX2 in breast cancer tissues. We collected 42 pairs of freshly frozen ERα-positive breast cancer tissues and matched adjacent noncancerous tissues. Western blot analysis revealed that MSX2 protein levels were markedly lower in tumor tissues compared with the adjacent controls (Figure 2A-B). Consistently, qPCR analysis of 12 paired of samples showed that MSX2 mRNA expression was reduced in tumors (Figure 2C). To evaluate the clinical relevance of MSX2, we further performed immunohistochemical (IHC) staining on commercial tissue microarrays and paraffin sections with detailed patient information. These results demonstrated a gradual decrease in MSX2 expression with increasing tumor grade (Figure 2D-E). Univariate analysis indicated that MSX2 expression correlated significantly with histological grade, pathological stage, and Ki67 index, but not with other clinicopathological parameters (Table 1). Kaplan–Meier survival analysis revealed that higher MSX2 expression predicted better overall survival (Figure 2F), suggesting its potential as a favorable prognostic biomarker in breast cancer.

**Figure 2.**
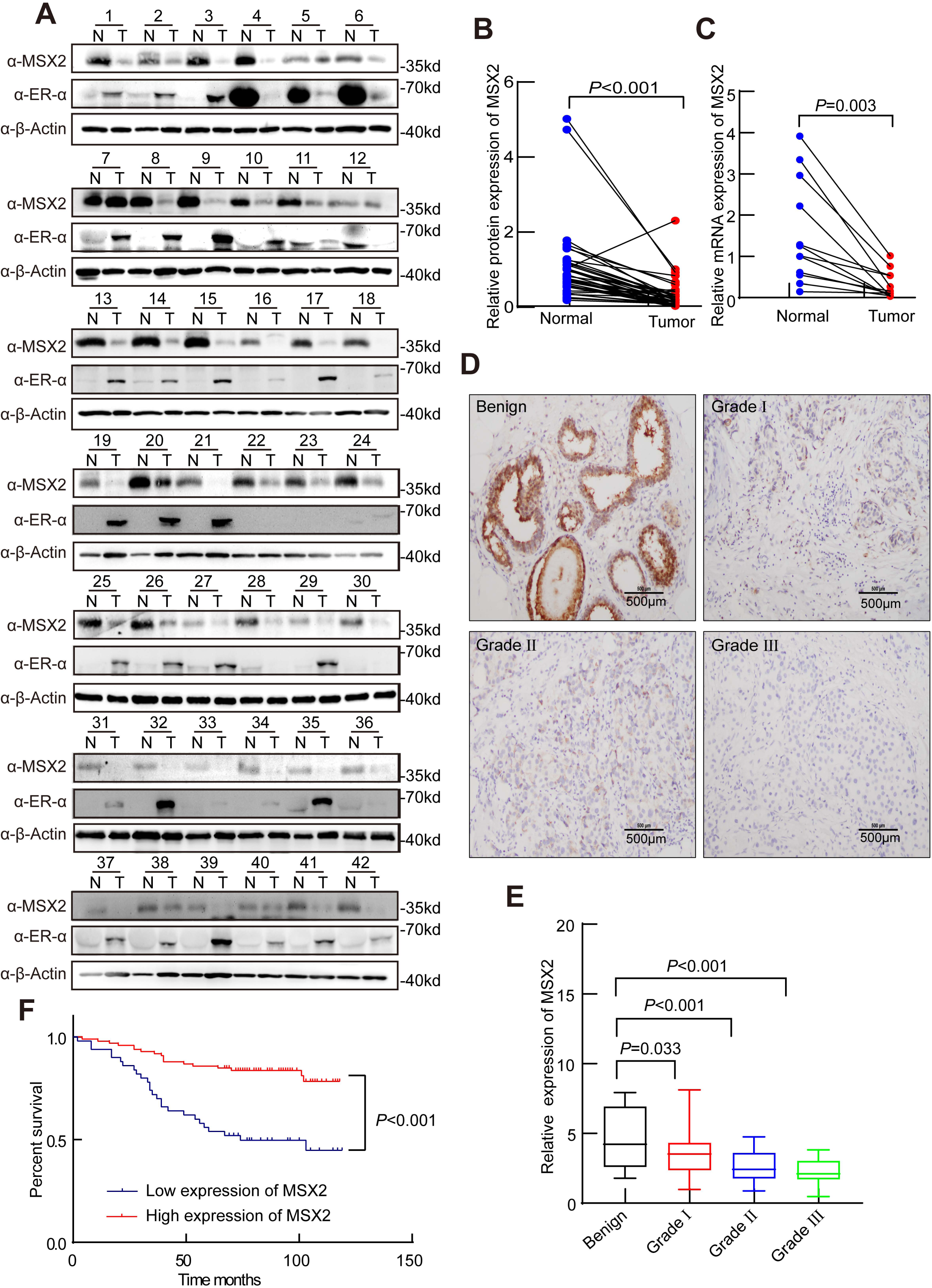
MSX2 is lower expressed in clinical breast cancer samples. **A:** The protein expression of MSX2 and ERα in 42 presentative pairs of primary BCa (T) and adjacent non-cancerous tissues (N). **B:** Grayscale analysis of MSX2 level was conducted using β-actin as the internal control and the differential expression of MSX2 between tumor tissues (T) and adjacent non-cancerous tissues (N) was delineated. \*\*\**p* < 0.001 (mean ± SD; Student’s t test; n = 42.) **C:** qPCR analysis showing the relative mRNA level of MSX2 in fresh human breast tissues. \*\**p* < 0.01 (mean ± SD; Student’s t test; n = 12). **D:** Representative MSX2 IHC images of primary ERα-positive breast tumors in different grades. The scale bars represent 500 µm. **E:** Statistical quantifications of MSX2 expression in benign tissues (n = 13) and different grades of breast cancer tissues (grade I n = 67, grade II n = 36, grade III n = 27). The central mark indicates the median, and the bottom and top edges of the box indicate the 25th and 75th percentiles, respectively. \*\*\**p*< 0.001 and \**p* < 0.05 (mean ± SD; Student’s t test). **F:** A higher MSX2 level predicts a favorable clinical outcome in breast cancer (BCa) cases (n = 149). The median expression score of MSX2 was used as the cutoff to assess overall survival using the Kaplan–Meier (KM) method. \*\*\**p* < 0.001 (mean ± SD; Student’s t test).

**Table 1:** Relationship between expression of MSX2 and clinical pathologic features in breast cancer.

| Characteristics | Cases(N=149) | MSX2 expression |  | P-value |
| --- | --- | --- | --- | --- |
|  |  | Low<br>(N=50) | High<br>(N=99) |  |
| Age |  |  |  |  |
| ≤55 | 67 | 28 | 39 | 0.054 |
| >55 | 82 | 22 | 60 |  |
| T |  |  |  |  |
| T0/T1 | 39 | 7 | 32 | 0.024 |
| T2+ | 106 | 40 | 66 |  |
| missing data | 4 | 3 | 1 |  |
| N |  |  |  |  |
| N0/N1 | 115 | 39 | 76 | 0.777 |
| N2+ | 32 | 10 | 22 |  |
| missing data | 2 | 1 | 1 |  |
| Histological grade |  |  |  |  |
| I-II | 73 | 17 | 56 | 0.012 |
| II-III | 40 | 17 | 23 |  |
| III | 27 | 14 | 13 |  |
| missing data | 9 | 2 | 7 |  |
| ERα status |  |  |  |  |
| ERα- | 62 | 25 | 37 | 0.122 |
| ERα+ | 82 | 23 | 59 |  |
| missing data | 5 | 5 | 0 |  |
| PR status |  |  |  |  |
| PR- | 71 | 37 | 60 | 0.092 |
| PR+ | 58 | 11 | 35 |  |
| missing data | 6 | 2 | 4 |  |
| HER2 status |  |  |  |  |
| HER2- | 102 | 37 | 65 | 0.255 |
| HER2+ | 45 | 12 | 33 |  |
| missing data | 2 | 1 | 1 |  |
| ki67 status |  |  |  |  |
| ki67- | 66 | 16 | 50 | 0.033 |
| ki67+ | 78 | 32 | 46 |  |
| missing data | 5 | 2 | 3 |  |
| AR status |  |  |  |  |
| AR- | 44 | 19 | 25 | 0.098 |
| AR+ | 103 | 30 | 73 |  |
| missing data | 2 | 1 | 1 |  |

### MSX2 interacts with ERα in breast cancer cells

Clinical diagnostic data show that more than 60% to 70% of breast cancer patients are ER-positive, and more than 80% of positive patients have high expression of ERα(Giaquinto *et al*, 2024). Having shown that MSX2 is weakly expressed in ER-positive tumor tissues, suggesting that MSX2 may inhibit breast cancer development. Additionally, data analysis revealed a negative correlation between MSX2 and the ERα downstream target genes MYC and E2F1. Therefore, we then asked whether MSX2 functions as a co-factor of ERα. First, STRING database (https://cn.string-db.org/) suggested that MSX2 might correlate with ERα (Figure S1A). Then, we set out to examine the interaction between MSX2 and ERα in mammalian cells using Co-immunoprecipitation (Co-IP). The results showed that endogenous MSX2 can interact with the ERα in MCF-7 cells (Figure 3A-B). In addition, expression plasmids for ERα and FLAG-MSX2 were co-transfected into HEK293 cells. Specific antibodies targeting either FLAG or ERα were utilized to precipitate the corresponding proteins as indicated. Our findings demonstrated that MSX2 interacts with ERα (Figure 3C-D).Using glutathione-S-transferase (GST) pull-down assay was performed with GST-tagged ERα-AF1 (29-180aa) or ERα-AF2 (282-595aa) fragments(Sun *et al*., 2020), we found that MSX2 directly binds ERα-AF1 fragment in vitro (Figure 3E). Additionally, we performed immunofluorescence (IF) experiments to evaluate the subcellular localization of MSX2 and ERα within MCF-7 and T47D cells. The findings indicated that MSX2 was present in the nucleus regardless of E2 treatment, whereas ERα diffused into the nucleus and entirely co-located with MSX2 in the presence of E2(Figure 3F-G). Overall, these results imply that MSX2 physically associates with ERα in mammalian cells.

**Figure 3.**
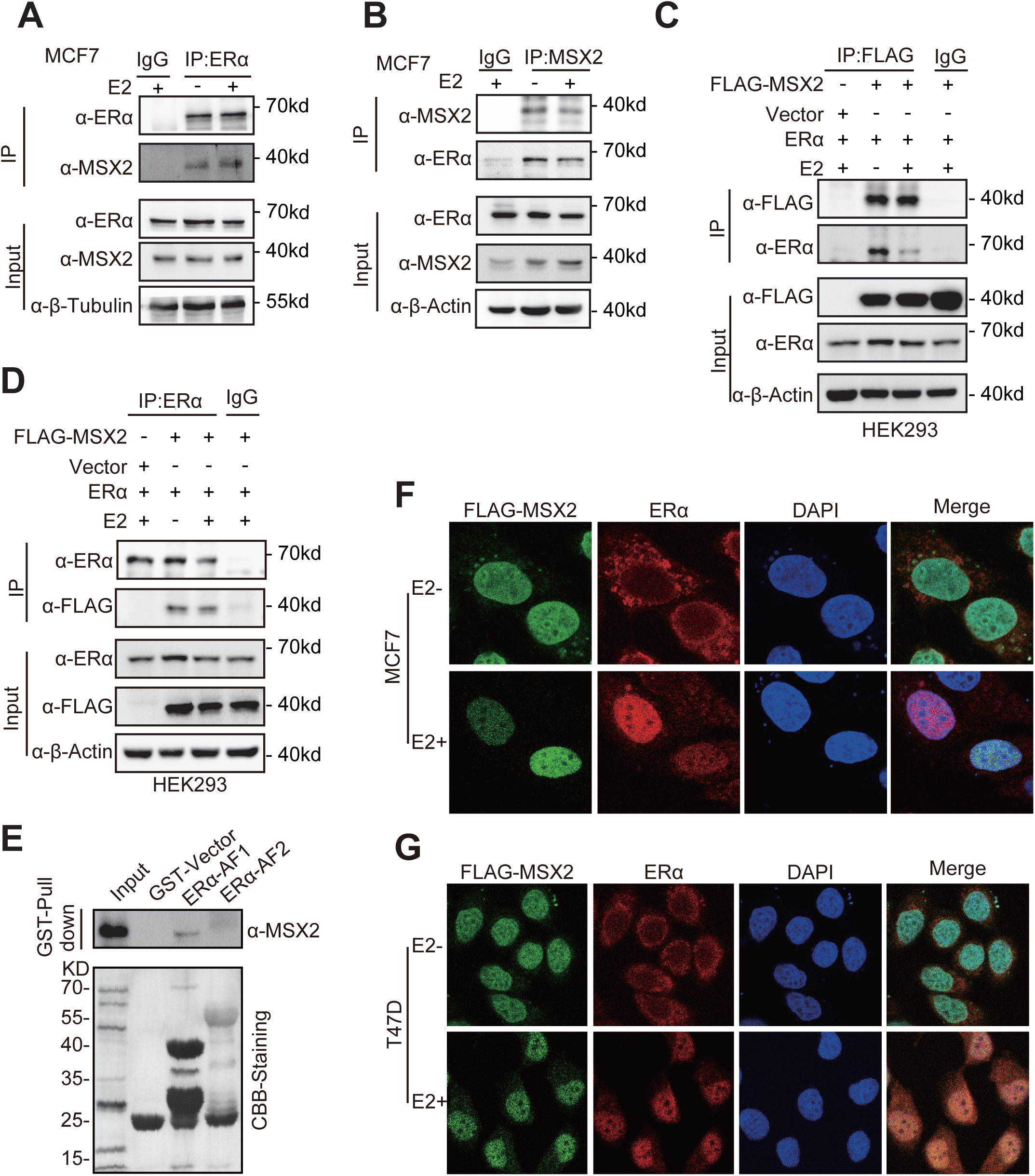
MSX2 interacts with ERα in ERα-positive breast cancer cells. **A-B:** Co-IP was utilized to examine the interaction between endogenous MSX2 and ERα in MCF-7 cells. Immunoprecipitation was conducted using anti-ERα and anti-MSX2 antibodies as primary antibodies, respectively. **C-D:** Additionally, Co-immunoprecipitation (Co-IP) assays confirm the interaction between exogenous MSX2 and ERα. FLAG-MSX2 and ERα expression plasmids were co-transfected into HEK293 cells, which were incubated in phenol red-free medium for 48 hours and subsequently treated with E2 (100 nM) for 12 hours. Whole cell extracts were collected for immunoprecipitation using anti-FLAG or IgG antibodies (C) or using anti-ERα or IgG antibodies (D), with 5% retained as an input control. **E:** The direct interaction between MSX2 and ERα was investigated using a GST-pull down assay. Proteins, including GST, GST ERα-AF1, and GST ERα-AF2, produced in an *E. coli* system, were combined with in vitro synthesized 35S-MSX2 protein. The proteins that bound during the reactions were analyzed using SDS-PAGE and autoradiography, with asterisks indicating the positions of the GST fusion proteins. **F-G:** Expression plasmids for ERα and FLAG-MSX2 were co-transfected into MCF-7 and T47D cells. Immunofluorescence analysis reveals the co-localization of Flag-MSX2 (green) and ERα (red) in MCF-7 (F) and T47D (G) cells. The nuclei were stained with DAPI (blue), and scale bars represent 10 µm.

### MSX2 represses ERα-mediated transactivation in mammalian cells

Having established that MSX2 interacts with ERα, we sought to investigate its role in the ERα-mediated regulation of gene transcription. To this end, we conducted luciferase reporter assays in MCF-7 and HEK293 cells. The results demonstrated that MSX2 significantly downregulated ERα-mediated transcription activity in the presence of estrogen (Figure 4A-B). To elucidate which functional domains of ERα are regulated by MSX2, we performed luciferase reporter assays in HEK293 (Figure 4C) and MCF-7 cells (Figure S1B), respectively. The results indicated that both the ERα-AF1 and ERα-AF2 domains contribute to the downregulation of ERα by MSX2, regardless of the presence of estrogen. These findings suggest that MSX2 may serve as a novel coregulator of ERα.

**Figure 4.**
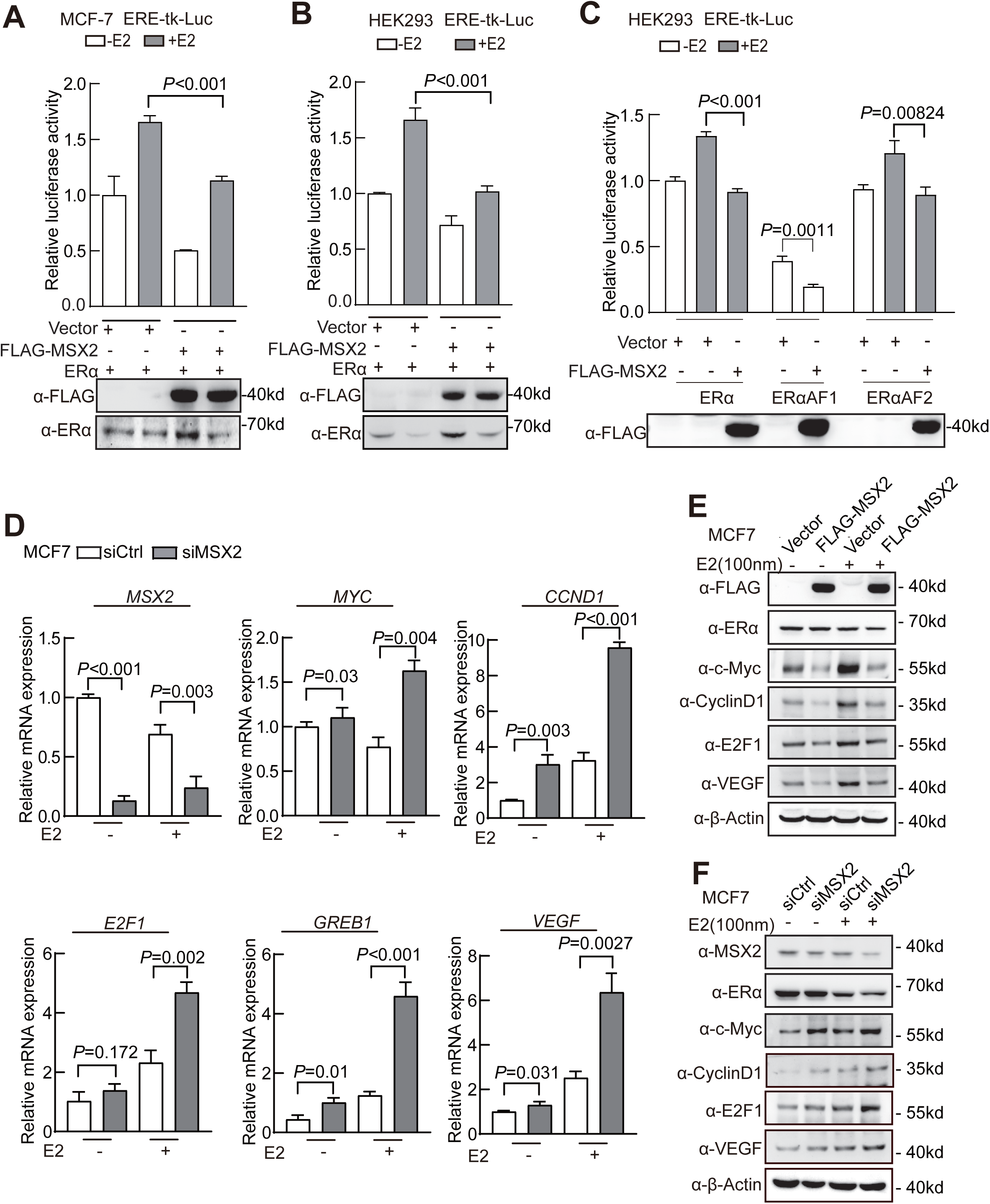
MSX2 inhibits ERα-mediated transactivation in mammalian cells. **A-B**: Relative luciferase activities were measured in HEK293 cells or MCF-7 cells that were transfected with the ERα expression plasmid, ERE-tk-luc, pRL-tk, and FLAG-tagged MSX2 expression plasmid (FLAG-MSX2) or PcDNA3.1 expression plasmid as Vector, both with and without the presence of E2 (100 nM).The expression levels of FLAG-MSX2 and ERα were detected using anti-FLAG and anti-ERα antibodies through western blotting. **C**: MSX2 downregulates ERα, ERα-AF-2 or ERα-AF-1 mediated transcriptional activity. HEK293 cells were transfected with either full-length ERα or truncated mutants containing ERα-AF-1 or ERα-AF-2, in the presence or absence of MSX2 expression. The expression levels of FLAG-MSX2 was detected using anti-FLAG antibodies through western blotting. **D**: qPCR analysis was conducted to assess the levels of transcript for verified ERα target genes in MCF-7 cells with MSX2-depleted. **E**: The effect of MSX2 depletion on estrogen-induced gene expression was assessed in MCF-7 cells transfected with control siRNA (siCtrl) or MSX2-specific siRNA (siMSX2). After treatment with 100 nM E2 for 16–18 hours, protein levels of ERα and its target genes were analyzed by Western blot. **F**: The impact of FLAG-MSX2 overexpression on estrogen-induced gene expression was examined by Western blot. MCF-7 cells were transfected with either Vector or FLAG-MSX2 plasmids and treated with or without 100 nM E2. The results presented above (A-F) represent three independent experiments that were conducted in duplicate. Data are presented as mean ± SD and analyzed using Student’s t-test. *p*-values are presented as exact values.

Indeed, ERα plays a crucial role in various biological processes through its target genes. To investigate the role of MSX2 in regulating endogenous ERα target genes, we conducted real-time quantitative PCR (qPCR) experiments. MSX2 was knocked down using MSX2-specific siRNA (siMSX2) in the MCF-7(Figure 4D) and T47D (Figure S1B) cell lines. The results indicated that the depletion of MSX2 led to a significant increase in the mRNA expression levels of ERα target genes, including *c-Myc, CyclinD1, E2F1, VEGF, GREB1 and TFF1* (Figure 4D and Figure S1C). We also examined the influence of MSX2 on the protein expression of a series of classical ERα target genes in MCF-7 and T47D cells. The results demonstrated that the ectopic expression of MSX2 resulted in a decrease in the protein levels of c-Myc, CyclinD1, and E2F1 (Figure 4E and Figure S1D). Meanwhile, the depletion of MSX2 increased the protein levels of ERα-regulated genes, such as c-Myc, CCND1, and E2F1 (Figure 3F and Figure S1E). These experimental results indicate that MSX2 functions as a corepressor of ERα, downregulating ERα-mediated gene transcription and modulating multiple downstream target genes of ERα.

### MSX2 is recruited to the *cis*-regulatory element regions of ERα target genes

To further investigate the molecular mechanism of modulation function of MSX2 on ERα action, we performed chromatin immunoprecipitation (ChIP) assays in two ERα-positive breast cancer cell lines, MCF-7 and T47D. We examined the recruitment of MSX2 and ERα to the ERE upstream of the transcription start site (TSS) of *c-Myc*, a well-established ERα target gene(Sun *et al*., 2020).The results demonstrated that both MSX2 and ERα were recruited to the ERE region of *c-Myc* in MCF-7 cells (Figure 5A) and T47D cells (Figure 5B). Additionally, ChIP assays were performed to evaluate the recruitment of MSX2 and ERα to the EREs of several ERα target genes, including *c-Myc, CyclinD1, E2F1, VEGF, GREB1, and TFF1*. We observed that ERα enrichment at the ERE regions of these genes was significantly increased in MSX2-depleted MCF-7 cells upon E2 treatment (Figure 5C) and showed a consistent pattern in T47D cells (Figure S2A). To determine whether MSX2 and ERα are co-recruited to the c-Myc promoter, ChIP-re-ChIP assays were performed. The results showed that MSX2 and ERα were simultaneously recruited to the ERE region of the c-Myc promoter in the presence of E2 in both MCF-7 (Figure 5D-E) and T47D cells (Figure S2B-C). Collectively, these findings indicate that MSX2 and ERα are recruited to the promoter regions of ERα target genes. Interestingly, MSX2 depletion enhances ERα recruitment to the ERE regions of ERα target genes, suggesting that MSX2 acting as a transcription factor may impede ERα recruitment to the promoter regions of ERα target genes in ERα-positive breast cancer cells.

**Figure 5.**
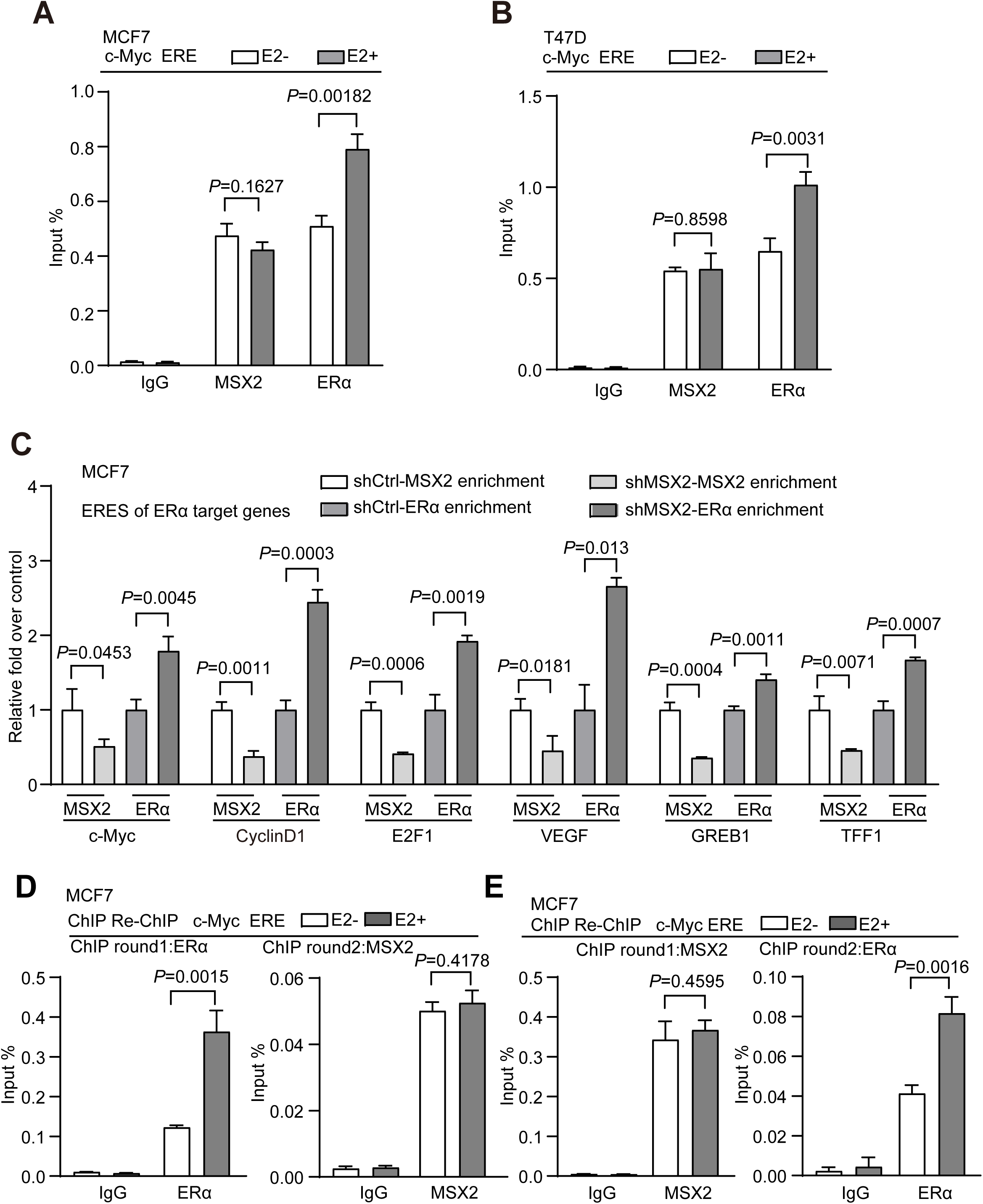
MSX2 and ERα are recruited to the promoter regions of ERα target genes. **A-B:** ChIP experiments were conducted to investigate the recruitment of MSX2 and ERα to the c-Myc promoter region in MCF-7 and T47D cells, both in the presence and absence of estrogen (100 nM). **C:** ChIP analysis of MSX2 and ERα was performed in MSX2-deficient MCF-7 cells, focusing on specific ERα-binding sites of E2-induced genes as indicated. The cells were cultured in estrogen-depleted medium for 48 hours prior to stimulation with 100 nM E2 for 2 hours. The immunoprecipitated DNA fragments were subsequently analyzed by qPCR using primers that target the promoter regions of ERα target genes. The amplified products were normalized against a defined quantity of unprecipitated input DNA. **D-E:** ChIP-re-ChIP experiments confirmed that E2 treatment enhanced the recruitment of ERα to the c-Myc-ERE in MCF-7 cells, while the recruitment of MSX2 remained largely unchanged. Chromatin extracts were immunoprecipitated using anti-ERα(D) or anti-MSX2(E) as the primary antibodies, followed by re-immunoprecipitation with anti-MSX2 (D) or anti-ERα (E), respectively. The immunoprecipitated DNA was then analyzed by qPCR to quantify the DNA signals. The results shown in panels (A-E) represent three independent experiments. *p*-values are provided as exact values. Data are presented as mean ± SD and were analyzed using Student’s t-test.

### MSX2 associates with HDAC1/HDAC2 complex to suppress ERα-mediated transcription

Our study has shown that MSX2 is recruited to the ERE regions of ERα target genes and suppresses transcription. Given the critical role of chromatin structure and histone modifications in the regulation of gene transcription, we hypothesize that MSX2 may further enhance its transcriptional repression by recruiting chromatin-modifying enzymes. STRING protein network analysis suggests that MSX2 associates with ERα and HDAC1/HDAC2 (Figure S3A). While HDAC1 and HDAC2 have been implicated in various aspects of transcriptional regulation and physiological processes(Asmamaw *et al*, 2024; Kelly & Cowley, 2013), their potential role in ERα-mediated gene transcription in conjunction with MSX2 remains poorly understood. Therefore, to investigate the potential role of MSX2 in modulating the action of HDAC1/HDAC2 during ERα-mediated transcription, we performed a series of experiments. Co-IP assays showed that MSX2 interacted with HDAC1 and HDAC2 (Figure 6A and Figure S3B). In addition, the interaction between ERα and HDAC1/HDAC2 had no significant changes when MSX2 is depleted (Figure 6B and Figure S3C), suggesting that MSX2 is not essential for the interaction between ERα and HDAC1/HDAC2, but may influence the functional association between these proteins.

**Figure 6.**
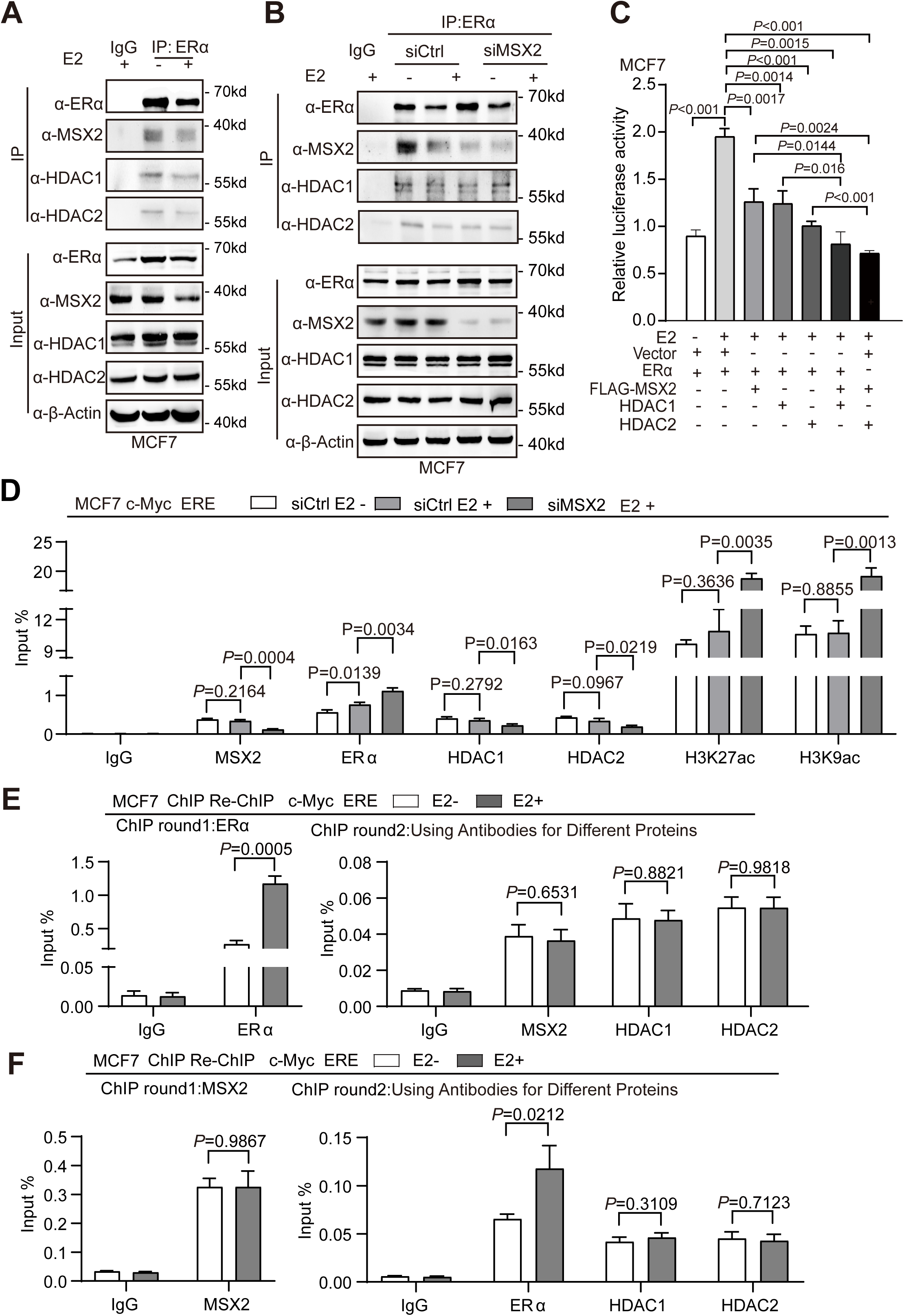
MSX2 is required for the recruitment of HDAC1/HDAC2 complex to the promoter regions of ERα target genes. **A:** Co-immunoprecipitation (Co-IP) was performed to investigate the interaction between endogenous MSX2, ERα, and HDAC1/HDAC2 proteins in MCF7 cells. The immunoprecipitation was carried out using anti-ERα antibodies to pull down MSX2-associated proteins, followed by the detection of ERα, HDAC1, and HDAC2 as potential interacting partners. **B:** Co-IP experiments were performed in MCF7 cells transfected with either control siRNA (siCtrl) or siMSX2, as indicated. Immunoprecipitation was carried out using anti-ERα and IgG antibodies, and the precipitated proteins were analyzed by western blotting with antibodies against HDAC1/HDAC2 or MSX2/ERα, as specified. **C:** The cooperative effect of MSX2 and HDAC1/HDAC2 proteins on the co-repression of ERα-mediated transactivation was evaluated. MCF-7 cells were co-transfected with ERα along with FLAG-MSX2, HDAC1, or HDAC2 expression plasmids as indicated for luciferase reporter assays. Vector plasmids were used as the control. **D:** ChIP assays were performed using specific antibodies to investigate the effect of MSX2 knockdown on the recruitment of ERα and HDAC1/HDAC2 proteins, as well as the corresponding histone modifications at the c-Myc ERE in MCF-7 cells. **E-F:** The recruitment of ERα (E) or MSX2 (F) together with each component of HDAC1/HDAC2 proteins on c-Myc-ERE by ChIP re-ChIP assays in MCF-7 cells. The results shown in panels (C-F) are representative of three independent experiments. *p*-values are presented as exact values. Data are expressed as mean ± SD and were analyzed using Student’s t-test.

To assess the functional consequences of these interactions, we employed luciferase reporter assays. Luciferase assays results showed that MSX2 or HDAC1/HDAC2 proteins respectively inhibits ER-mediated transactivation, and MSX2/HDAC1, MSX2/HDAC2L could additionally co-repress ERα action in MCF-7 cells (Figure 6C). Furthermore, ChIP assays were performed in MCF-7 cells. The results demonstrated that MSX2, ERα, and HDAC1/HDAC2 were recruited to the promoter region of the *c-Myc*. Additionally, the depletion of MSX2 led to an increased recruitment of ERα and a decreased recruitment of HDAC1 and HDAC2, which was accompanied by elevated levels of histone H3K27ac and H3K9ac levels (Figure 6D). To further confirm the role of MSX2 in regulating the recruitment of HDAC1/HDAC2 and its impact on histone modifications, we performed additional ChIP assays in MCF-7 cells ectopically expressing MSX2. ChIP assay results showed that ectopic expression of MSX2 could respectively reduce the recruitment of ERα, while increase the recruitment of HDAC1 and HDAC2, which subsequently reduced the levels of H3K27ac and H3K9ac (Figure S3D). ChIP re-IP assays were further performed to explore whether ERα and MSX2 could be recruited together with HDAC1/HDAC2 complex to the promoter region of the ERα target gene. The results showed that ERα and MSX2 were recruited to MYC-ERE region together with HDAC1/HDAC2. Notably, the recruitment of ERα is enhanced in the presence of estrogen, whereas the recruitment of MSX2 and HDAC1/HDAC2 remains unaffected (Figure 6E-F and Figure S3E-F). The results indicate that MSX2 interacts with HDAC1 and HDAC2 to collaboratively inhibit the recruitment of ERα to the promoter region of its target genes, thereby altering the modification levels of H3K27ac and H3K9ac. Taken together, our results suggest that MSX2 associates with ERα and HDAC1/HDAC2 to inhibit ERα-mediated transcriptional activity in ERα-positive breast cancer cells.

### MSX2 suppresses cell growth in ERα-positive breast cancer cells

Having established the molecular mechanisms by which MSX2 influences ERα action, we then proceeded to investigate the potential biological role of MSX2 in the progression of ERα-positive breast cancer. We established the MCF-7 and T47D cell lines with stable ectopic expression or knockdown of MSX2 (OEMSX2 and shMSX2) with induced by lentiviral infection, the efficiency of MSX2 overexpression and knockdown was verified by Western blot analysis (Figure S4A-B). Growth curve analysis revealed that MSX2 overexpression significantly inhibited cell proliferation in both MCF-7 (Figure 7A) and T47D cells (Figure S4C), whereas knockdown of MSX2 promoted cell growth (Figure 7B and Figure S4D). Consistently, colony formation assays demonstrated that MSX2 overexpression markedly suppressed colony formation (Figure 7C and Figure S4E), while MSX2 knockdown enhanced colony-forming ability (Figure 7D and Figure S4F). To further investigate its function, we performed flow cytometry to evaluate the effect of MSX2 on cell cycle progression. The results showed that MSX2 overexpression delayed the G1-S phase transition, causing G1 phase arrest (Figure 7E and Figure S4G), while MSX2 knockdown promoted cell cycle progression (Figure 7F). Collectively these findings suggest that MSX2 functions as a tumor suppressor by inhibiting cell proliferation and blocking the G1-S phase transition in ERα-positive breast cancer.

**Figure 7.**
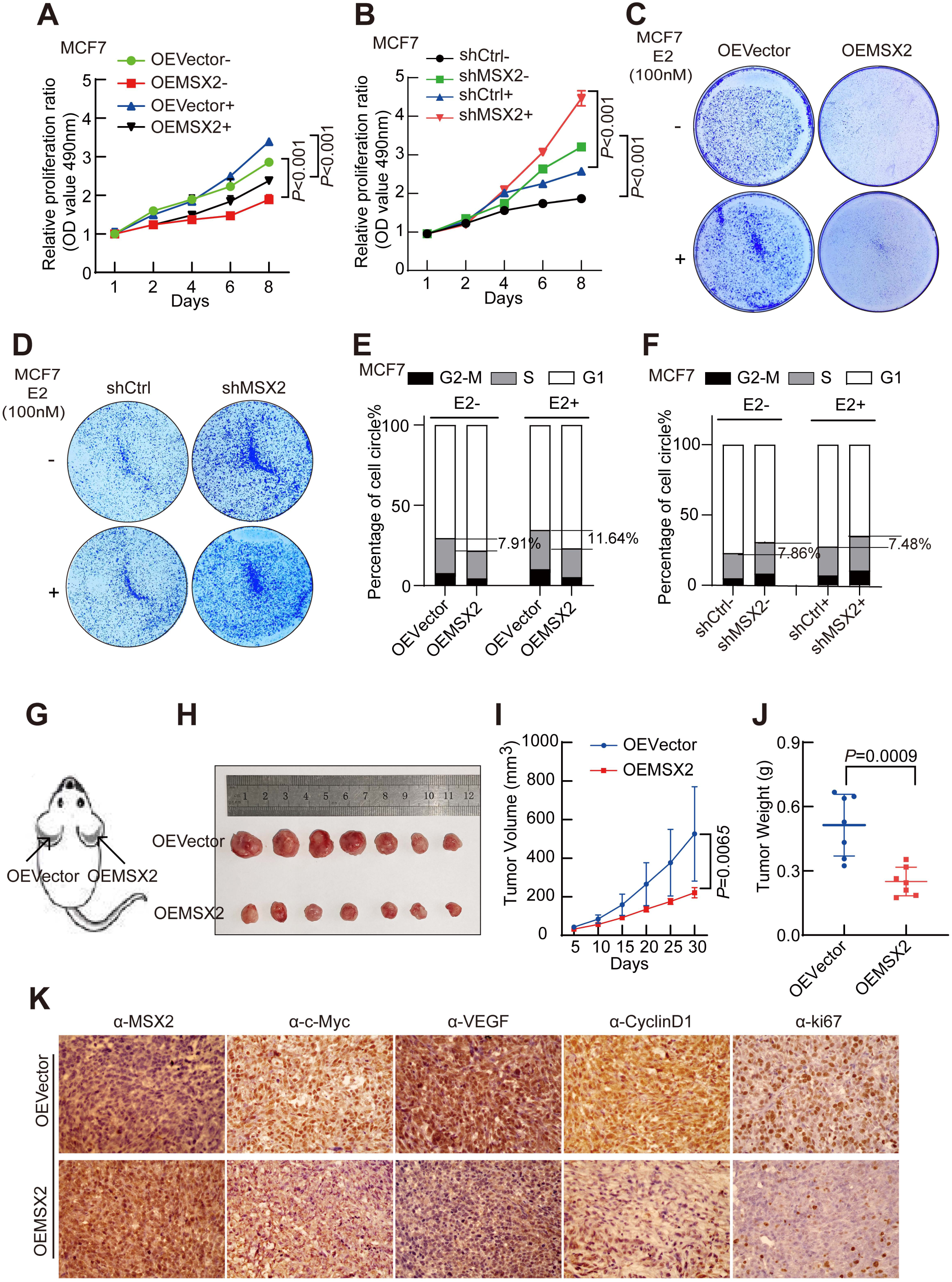
MSX2 inhibits cell growth in ERα-positive breast cancer cells. **A-B:** Growth curves showing the effect of MSX2 overexpression (A) or knockdown (B) on MCF-7 cells proliferation in the presence or absence of E2 (100 nM). Total cell viability was measured every other day using the MTS assay. **C-D:** The effects of MSX2 on cell growth in MCF-7 cells with OEMSX2 or shMSX2 were evaluated using a colony formation assay under vehicle or E2 (100 nM) treatment for 15 days. **E-F:** The representative histogram shows the cell cycle distribution of MCF-7 cells after transduction with OEMSX2(E) or shMSX2(F), illustrating the proportion of cells in the G0/G1, S, and G2/M phases under E2 stimulation or not, as determined by flow cytometry. All the above experiments were conducted using a scrambled OECtrl or shctrl as the control. **G-H:** Representative images depict the xenograft tumor in female BALB/c mice, which were injected with either the OECtrl (top left) or cells exhibiting OEMSX2 (bottom right), while maintaining a normal feeding pattern. **I**: The average tumor volume of the OECtrl and OEMSX2 groups was measured every five days, starting on the fifth day. **J:** Tumor weights were assessed between the two groups of tumors 30 days later. **K:** Paraffin sections of the tumors from nude mice were subjected to immunohistochemistry (IHC) staining using anti-MSX2, anti-c-Myc, anti-CyclinD1, anti-E2F1 and anti-Ki67 antibodies. Representative images were captured representing 200μm in the microscopic field. In all data, Student t-tests were used. Error bars represent mean ± SD. *p*-values are presented as exact values. \**p* < 0.05, \*\**p* < 0.01, \*\*\**p* < 1e−3 and *p* > 0.05 stands for no significance.

To investigate the in vivo effect of MSX2, we established an ectopic xenograft tumor model using MCF-7 cells that were infected with OEMSX2 or OEVector. 4-week-old female BALB/c nude mice were injected with the two types of infected cells in both axillary regions, and we monitored tumor growth and measured tumor sizes every five days post-injection. Compared to OEVector-MCF-7 cells, OEMSX2-MCF-7 cells exhibited reduced tumor growth rates (Figure 7G-H). In accordance with the growth curve, the tumor volumes from OEMSX2-MCF-7 cells exhibited a slower growth rate than those from the OEVector-MCF-7 cells, and the average tumor weight was also lower in the OEMSX2 group compared to OEVector (Figure 7I-J).These results prompted us to perform quantitative RT-PCR to examine the expression changes of estrogen-responsive genes. As shown in Figure S4H-L, MSX2 overexpression significantly reduced the mRNA levels of *c-Myc*, *CyclinD1, E2F1* and *VEGF* in xenograft tumor tissues. In addition, immunohistochemical (IHC) analysis was performed to examine the expression in MSX2 and target expression of ERα. The results revealed that ectopic expression of MSX2 significantly reduced the expression levels of c-Myc, CyclinD1, E2F1, and VEGF, accompanied by a reduction in the Ki67 index in xenograft tumor tissue (Figure 7K). Collectively, these findings indicate that the ectopic expression of MSX2 inhibits breast cancer cell growth in mice and reduces the expression of estrogen-induced genes in vivo.

### MSX2 facilitates the sensitivity of ERα-positive breast cancer cells to antiestrogen treatment

Generally, estrogen inhibitors or ERα antagonists, which function through the estrogen-ERα axis, have been the predominant treatment for ERα-positive breast cancer, including aromatase inhibitors (such as Letrozole), selective ERα modulators (such as Tamoxifen) and selective ERα degraders (such as Fulvestrant). Having established MSX2 participates in modulation of ERα signaling pathway, we further evaluated its role in ERα-positive breast cancer cell survival in response to the treatment of ERα inhibitors. We utilized T47D and MCF-7 cells stably overexpressing MSX2 and treated them with an ERα antagonist, followed by colony formation assays to assess cell proliferation. MSX2 overexpression in combination with appropriate concentrations of tamoxifen, fulvestrant, or letrozole, resulted in increased cell death compared to the group treated only with E2 (Figure 8A and Figure S5A). Furthermore, the drug treatment cell growth curves demonstrated comparable results in T47D and MCF-7 cells at the appropriate drug concentrations (Figure 8B-E and Figure S5B-E). Collectively, these findings suggest that MSX2 may be linked to the sensitivity of endocrine drugs.

**Figure 8.**
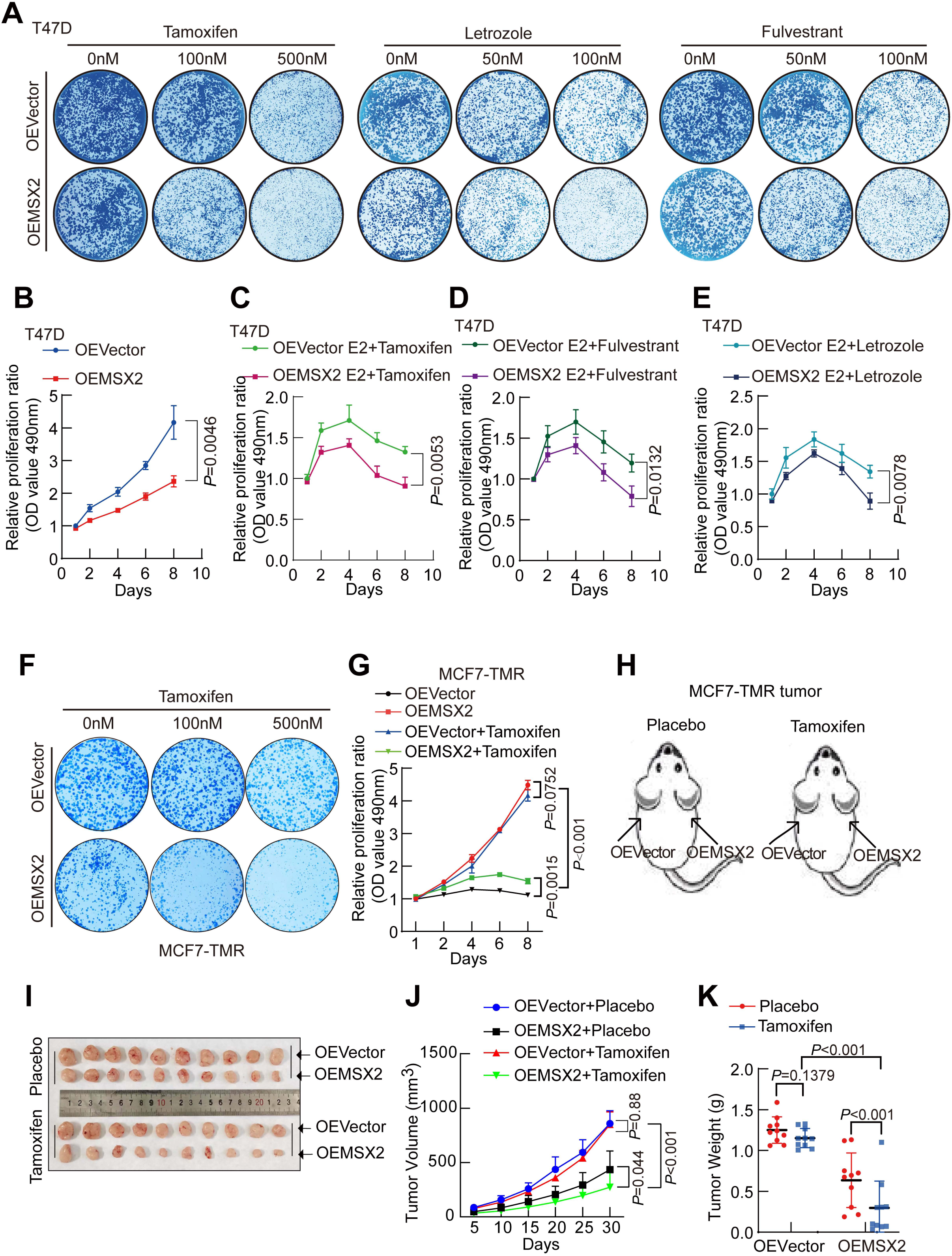
MSX2 enhances the sensitivity of anti-estrogen treatment in ERα-positive breast cancer cells. **A:** The influence of MSX2 on endocrine drug sensitivity in T47D cells was investigated. T47D cells were pre-transfected with OEMSX2 or OECtrl. The cells were cultured in varying concentrations of three types of endocrine drugs for a duration of 15 days. Subsequently, the cells in the wells were stained with R250, and the photographs presented represent one of three independent experiments. **B-E:** The growth curve illustrates the effect of MSX2 overexpression on the proliferation of T47D cells under different conditions: in the absence of endocrine drugs (B), and in the presence of Tamoxifen (500 nM) (C), Fulvestrant (100 nM) (D), and Letrozole (100 nM) (E). Total cell viability was measured every other day using the MTS assay. **F:** The panels show the results of colony formation assays conducted in MCF-7TRM cells. The cells were infected with lentiviruses expressing either OECtrl or OEMSX2. In each panel, the cells were treated with different doses of Tamoxifen for 15 days, followed by fixation and staining with Coomassie Brilliant Blue R-250. **G:** Growth curve showing the effect of MSX2 on MCF-7 TMR cells proliferation with or without Tamoxifen (1uM). Total cell viability was assessed every other day by MTS assay. **H-I:** Representative photographs of xenograft tumors in female BALB/c mice injected with OECtrl (left/up) and OEMSX2 (right/down) stable MCF-7 TMR cells. The vector cohort was fed with placebo tablets, while the experimental cohort received tamoxifen citrate. **J:** Tumor volume among the four groups demonstrated the effect of MSX2 on the intrinsic Tamoxifen sensitivity of MCF-7 TMR cells. Mice were administered either a placebo or tamoxifen citrate every three days. The drugs were applied on the fifth day, and the average tumor volume was measured every five days, beginning on the fifth day. The plot illustrates the tumor volumes. **K:** Data illustrating the tumor weight of mice after a 30-day period is presented. In all analyses, Student’s t-tests were employed. Error bars represent the mean ± standard deviation (SD). *p*-values are presented as exact values. \**p* < 0.05, \*\**p* < 0.01, \*\*\**p* < 1e−3 and *p* > 0.05 stands for no significance.

We also conducted relevant experiments using BT474 breast cancer cells, which are intrinsically resistant to tamoxifen, and Tamoxifen-resistant MCF-7 (MCF-7 TMR). MSX2 overexpression was induced in MCF-7 TMR and BT474 cells through lentiviral infection, and the efficiency of MSX2 overexpression was confirmed by Western blot analysis (Figure S5F). Colony formation assays demonstrated that tamoxifen alone had minimal effect on cell viability; however, the combined application of tamoxifen and MSX2 overexpression significantly inhibited colony formation in MCF-7 TMR (Figure 8F) and BT474 cells (Figure S5G). Similarly, MTS assays showed that MSX2 overexpression enhanced the sensitivity of MCF-7 TMR and BT474 cells to tamoxifen treatment (Figure 8G and Figure S5H). In summary, our findings suggest that MSX2 may play a role in modulating endocrine sensitivity in ERα-positive breast cancer.

Building on the previous findings regarding the biological role of MSX2 in tamoxifen resistance in cultured cells, we aimed to investigate the impact of MSX2 on tamoxifen treatment in vivo. MCF-7 TMR cells, which were transfected with ectopic expression of MSX2 through lentiviral infection (OEMSX2), along with vector control cells (OEVector), were injected into 4-week-old BALB/c Nude mice. As shown in Figure 8H-I, during the tumor growth process, each mouse was administered either tamoxifen citrate or placebo pellets to ensure consistent tamoxifen treatment. Our results demonstrated that ectopic expression of MSX2 could retard tumor growth, regardless of tamoxifen treatment. Compared to placebo treatment, tamoxifen treatment only modestly delayed tumor growth (Figure 8J); however, OEMSX2 significantly enhanced the tamoxifen effect, leading to increased cell death in MCF-7 TMR cells (Figure 8K). Consistent with the in vitro data, these findings indicate that MSX2 is associated with the sensitivity of antiestrogen treatments in ERα-positive breast cancer cells.

## Discussion

ERα-positive breast cancer represents a biologically diverse disease entity with spatiotemporal heterogeneity, variable therapeutic responses, and propensity for drug resistance. Understanding this heterogeneity is pivotal to unlocking truly individualized precision medicine to improve the patient survival (Andrade de Oliveira *et al*, 2023; Guo *et al*, 2023; Sivasailam *et al*, 2025). Decoding the multifaceted modulation of Erα signaling pathway is of considerable interest for finding the potential strategy for endocrine resistance in luminal subtype. In this study, we identified MSX2 as a novel ERα co-repressor in breast cancer. Downregulation of MSX2 is correlated with the poor prognosis in ERα-positive breast cancer. Our results have demonstrated that MSX2 associates with HDAC1/2 complex to inhibit ERα action, and MSX2 is required for the recruitment of HDAC1/2 complex to the promoter regions of ERα target genes. Functionally, our data indicate that MSX2 suppresses breast cancer-derived cell growth/proliferation and enhances the sensitivity to antiestrogen treatment in ERα-positive breast cancer (Figure 9).

**Figure 9.**
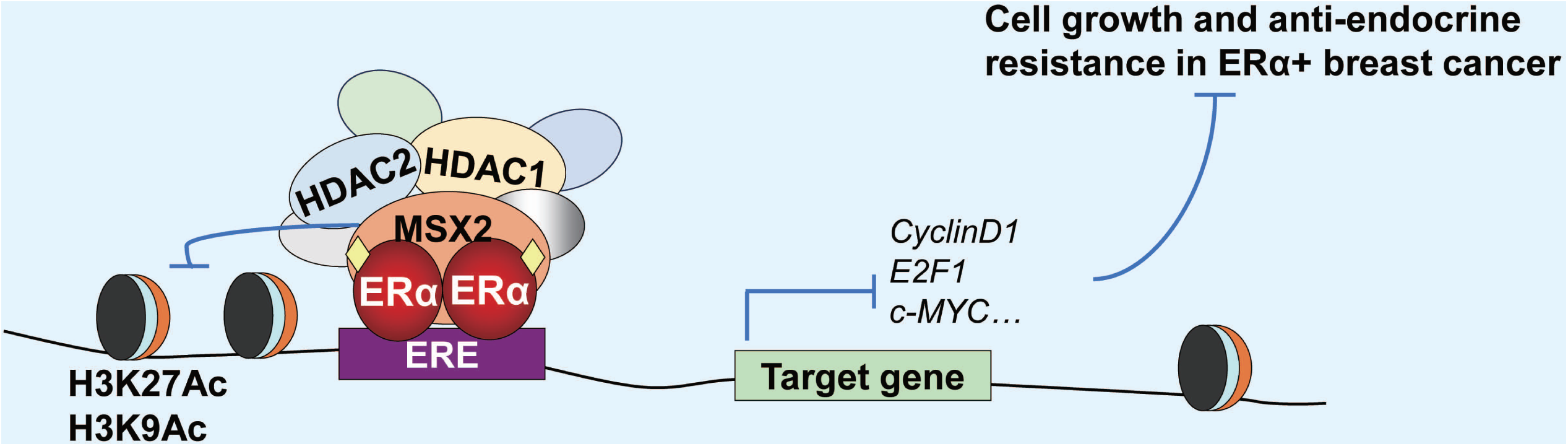
Schematic illustration for modulation function of MSX2 on ERα-mediated transactivation. Downregulation of MSX2 in breast cancer is correlated with the poor prognosis. MSX2 as a novel ERα co-repressor associates with HDAC1/2 complex. MSX2 is required for the recruitment of HDAC1/2 complex to the promoter regions of ERα target genes, reducing histone H3K27ac and H3K9ac levels to inhibit ERα-induced transactivation. MSX2 suppresses breast cancer-derived cell growth/proliferation and enhances the sensitivity to antiestrogen treatment in ERα-positive breast cancer.

Accumulating evidences indicate that MSX2 plays a complex role in tumorigenesis, progression, metastasis, and treatment response. Our data demonstrate that lower expression of MSX2 is significantly associated with poor survival in breast cancer patients (Figure 1 and 2). The marked downregulation of MSX2 in high-grade tumors, coupled with its strong correlation with poor survival, suggests its role as a gatekeeper of a differentiated, less aggressive state. This role aligns with the established function of homeobox genes in maintaining cellular differentiation of cancer and identity during tumor development. Consequently, the loss of MSX2 in advanced cancers may induce dedifferentiation at a developmental level, thereby relieving the suppression of ERα and enabling it to drive a more robust proliferative program.

This MSX2 loss-mediated liberation of ERα not only promotes proliferation but also has direct implications for treatment, given that the activity of ERα is the primary driver and therapeutic target in luminal breast cancer. Although the application of selective estrogen receptor modulators (SERMs, such as tamoxifen) has markedly improved patient survival, the development of endocrine resistance remains a major clinical challenge, limiting long-term benefits (Garcia-Martinez *et al*., 2021; Gong *et al*, 2017; Keikha *et al*, 2021; Yu *et al*, 2019a) .Given our finding that MSX2 loss liberates ERα, we hypothesized that MSX2 itself might be a critical regulator of endocrine therapy response. Therefore, we sought to determine whether MSX2 functions as a novel transcriptional repressor of ERα and whether its loss contributes to therapy resistance. Our study demonstrates that MSX2 is a potent repressor of ERα-mediated transcription. We found that MSX2 directly interacts with the AF1 domain of ERα via co-immunoprecipitation and GST pull-down assays (Figure 3). This interaction is particularly significant because the AF1 domain is crucial for ligand-independent activation, suggesting MSX2 may serve as a constitutive brake on ERα activity. Accordingly, luciferase assays showed that MSX2 significantly inhibits ERα-induced transactivation. Consistent with this, ectopic expression of MSX2 downregulated, while its deletion upregulated, key ERα target genes such as c-Myc, CyclinD1, and E2F1 (Figure 4 and S1). ChIP assay experiments confirmed that both MSX2 and ERα are recruited to estrogen response elements (EREs) of target genes (Figure 5 and S2). Intriguingly, MSX2 knockdown enhanced ERα binding to EREs, suggesting a competitive mechanism where MSX2 inhibits ERα recruitment or its transcriptional activity at these sites.

The regulation of ERα activity by co-regulators is a well-established mechanism of endocrine resistance. For instance, the corepressor NCoR(Bartella *et al*, 2012; Konduri *et al*, 2010; Wong *et al*, 2014), as a corepressor of ERα, interacts with the ligand-binding domain (LBD) of ERα, displacing coactivators such as the SRC family. NCoR also recruits HDACs that lead to chromatin condensation(Asmamaw *et al*., 2024; Keeton & Brown, 2005). Similarly, the tumor suppressor BRCA1 plays a crucial role in maintaining global heterochromatin integrity. Beyond its role in chromatin organization, BRCA1 directly interact with estrogen receptor alpha (ERα) to inhibit its transcriptional activity (Eakin *et al*, 2007; Mullan *et al*, 2006).In addition to BRCA1, other transcriptional repressors such as FOXC1 have been shown to negatively regulate ERα signaling. FOXC1 inhibits ERα expression and activity by blocking the interaction between GATA3 and the cis-regulatory elements (CREs) of the ERα gene. As a result, elevated FOXC1 expression attenuates the response of ER-positive breast cancer cells to both estrogen and tamoxifen treatment(Yu-Rice *et al*, 2016). Our findings now position MSX2 as a novel member of this group of ERα repressors. However, MSX2 is distinguished by its unique interaction with the AF1 domain—a key region for ligand-independent activation—and its identity as a homeobox protein. This directly links the regulation of ERα to fundamental developmental pathways. Consequently, the loss of this developmentally-programmed inhibitory mechanism may represent a pivotal event in the evolution of endocrine resistance.

Having established that histone deacetylases (HDACs) is involved in transcriptional repression within multiprotein complexes by altering chromatin structure(Asmamaw *et al*., 2024; Kelly *et al*, 2018; Seto & Yoshida, 2014). In this study, our results have demonstrated that MSX2 is involved in suppression of ERα-mediated transactivation. We then investigated a potential association between MSX2 and HDAC1/2 complex. We provided the evidences that MSX2 acts as functional partners of HDAC1/2. Moreover, MSX2 facilitates the recruitment of HDAC1/2 to estrogen response elements (EREs), resulting in decreased histone acetylation (H3K9ac and H3K27ac) (Figure 6 and Figure S3). This places MSX2 within a chromatin-modifying complex that fine-tunes ERα activity. Interestingly, MSX2 enhances HDAC1/2 recruitment but is not required for the ERα-HDAC interaction, suggesting MSX2 acts as a molecular guide that directs deacetylase activity to specific genomic loci for precision. This finding refines our understanding of how HDACs are recruited to the specific promoters to regulate gene transcription.

Our results also demonstrated that MSX2 inhibited cell proliferation even in the absence of E2, suggesting it may also suppress ligand-independent ERα signaling pathways, thereby restraining breast cancer progression (Figure 7 and Figure S4). Furthermore, the influence of MSX2 on the efficacy of antiestrogen treatment demonstrated that MSX2 overexpression increased the sensitivity of antiestrogen-resistant BCa cells to antiestrogen drug treatment (Figure 8 and Figure S5). From a translational perspective, our data strongly suggest that MSX2 is a critical determinant of endocrine therapy sensitivity. The synergy between MSX2 overexpression and tamoxifen or fulvestrant suggests that MSX2 and these drugs act on a common axis—ERα activity—through complementary mechanisms. While tamoxifen competes with estrogen for binding, and fulvestrant promotes degradation, MSX2 actively remodels chromatin to silence ERα targets. This multi-pronged attack is evidently more effective. Most importantly, the restoration of MSX2 sensitizing tamoxifen-resistant models (MCF-7 TMR and BT474) is a groundbreaking finding. It posits that the loss of MSX2 is not just a passive companion to progression but an active driver of resistance. This elevates MSX2 from a prognostic marker to a potential predictive biomarker and an independent attractive therapeutic target. Future studies exploring strategies to reactivate or mimic MSX2 function are warranted to determine their potential in reversing acquired endocrine resistance. Our work provides a strong rationale for exploring this exciting possibility.

In conclusion, our study identifies MSX2 as a critical negative regulator within the ERα transcriptional network, which functions by recruiting class I HDACs to fine-tune gene expression. This repressive activity not only deepens our mechanistic understanding of transcriptional control in breast cancer but also suggests novel therapeutic avenues. MSX2 is a substrate of the FBXW2 E3 ligase. The small-molecule inhibitor MLN4924 increases MSX2 expression by inactivating FBXW2, thereby sensitizing breast cancer cells to tamoxifen(Yin *et al*, 2019). Consistent with this model and our finding, MLN4924 represents a promising therapeutic candidate for ER-positive breast cancer. Future work should focus on delineating the full composition of the MSX2 repressor complex and validating the therapeutic efficacy of targeting this axis in vivo, with the therapeutic objective of overcoming therapeutic resistance in breast cancer patients.

## Materials and Methods

### Cell lines and cell culture

All cell lines were obtained from the American Type Culture Collection (ATCC) and authenticated through STR profiling. Prior to experimentation, they were tested and confirmed to be mycoplasma-negative. HEK293 (ATCC: CRL-1573) and MCF-7 (ATCC: HTB-22) were cultured in Dulbecco’s Modified Eagle Medium (DMEM) (Gibco), supplemented with 10% fetal bovine serum (FBS) (Gibco) and 100 U/ml penicillin-streptomycin (P/S). BT474 (ATCC: HTB-20) and T47D (ATCC: HTB-133) were cultured in RPMI-1640 medium (Gibco), also supplemented with 10% FBS and 100 U/ml P/S. When it was needed to add E2 stimulus, cells were grown in phenol red-free DMEM or RPMI-1640 containing 5% charcoal-treated serum. 17β-estradiol (E2, Sigma-Aldrich) was dissolved in ethanol (Aladdin). All cells were maintained in a humidified incubator at 37 °C with 5% CO2. The Tamoxifen-resistant MCF-7 (MCF-7 TMR) cell lines were cultured in phenol red-free DMEM containing 10% FBS and 1 μM Tamoxifen (Abmole, Cat# M7353).

### Plasmids and Antibodies

The MSX2 plasmid was sourced from SinoBiological. FLAG-tagged MSX2 was constructed by inserting the MSX2-FL plasmid into the pcDNA3 vector. ERα and ERE-Luc plasmids were kindly provided by Dr. Shigeaki Kato.

The antibodies were used in our study as follow: anti-MSX2 (santa,cat#sc-393986: Sigma, cat#HPA005652); anti-ER α (Cell Signaling Technology, cat# 8644); anti-FLAG (Shanghai Genomics, Cat# GNI4110-FG); anti-c-Myc (Proteintech, cat# 10828); anti-Cyclin D1 (Cell Signaling Technology, cat# 2978); anti-E2F1 (Proteintech, cat# 66515-1-Ig); anti-VEGF (Proteintech, cat# 19003-1-AP); anti-HDAC1 (Cell Signaling Technology, cat#34589); anti-HDAC2 (Cell Signaling Technology,cat#57156); anti-H3K9ac (Cell Signaling Technology, cat#9649); anti-H3K27ac (Cell Signaling Technology, cat#8173); anti-β -Actin (Proteintech,cat# 20536-1-AP); anti-GADPH (ABclonal, cat# AC036, 1:5000);anti-β -Tubulin (Proteintech,cat# 10094-1-AP); normal rabbit IgG (Santa Cruz, Cat# sc2027); Goat anti-mouse/rabbit IgG (H + L) secondary antibody, HRP (Invitrogen, Cat# 31430, 31460).

### Patients and tumor specimens

Human primary breast cancer (BCa) tissues and their corresponding adjacent normal tissues were obtained from Liaoning Cancer Hospital and the First Affiliated Hospital of China Medical University.

Ethical approval for this study was conducted by the Ethics Committee of the Research Department at China Medical University (no. 2025-032). The study was conducted in accordance with the ethical principles outlined in the Declaration of Helsinki. All participants were informed about the purpose of the study, assured of confidentiality, and provided written consent prior to participation. Participation was voluntary, and respondents could withdraw at any time without consequence.

### Western blotting and co-immunoprecipitation (co-IP) analysis

Western blotting was performed following the standard protocol previously described[43]. All experiments were performed with a minimum of three independent biological replicates.

For immunoprecipitation, experiments were conducted based on our previous study[24]. The whole cell lysates were extracted and equal protein amounts were immunoprecipitated with specific antibody or control IgG. And immune complexes were analyzed by western blot.

### Immunohistochemical (IHC) analysis

Breast tissue paraffin specimens were prepared at the First Hospital of China Medical University. The experimental procedure was based on our previous work [44], and this study was approved by the Ethics Committee of China Medical University. Commercial breast cancer tissue microarray slides (HBre-Duc150Sur-02) were obtained from Shanghai Outdo Biotech Co, Ltd (SOBC). Staining scores were evaluated using the H score method, which is based on the proportion of positively stained tumor cells (0–100%) and the intensity of brown staining (0–3). The final expression score, which ranged from 0 to 3 (negative = 0; weak = 1; moderate = 2; strong = 3), was calculated by multiplying these two independent indices.

### GST-pull down

GST fusion proteins GST ERα-AF1 (29-180 aa) and GST ERα-AF2 (282-595 aa) were puriffed by Glutathione-sepharose beads (GE Healthcare, Cat# 17075601). In vitro transcription and translation of the indicated proteins FLAG-MSX2 were performed by TNT-coupled transcription and translation system (Promega Corporation, Cat# L1171). The bound proteins were isolated by incubating the mixture for 2 h and washing three times, proteins were detected by western blot. Coomasssie brilliant blue stain indicated the loading amounts of the GST or GST-fusion proteins.

### Luciferase dual-reporter assays

Cells were subjected to serum starvation overnight and subsequently co-transfected with MSX2 expression plasmid, in addition to ERα, ERE-tk-Luc, and a control plasmid containing Renilla luciferase (pRL). E2 (100 nM) was then administered to the designated groups after 4 and 20 h, respectively, to maintain a continuous stimulation. Following 24 h of transfection, the cells were lysed in order to measure luciferase activity utilizing a Promega dual-luciferase reporter assay system (Promega Corporation, Cat# E1910). The average relative luciferase activity, as illustrated in the figure, is derived from at least three separate experiments.

### Immunofluorescence (IF)

Cells were fixed at room temperature using 4% paraformaldehyde for 20 min, permeabilized in PBS containing 0.05% Triton X-100 for an additional 20 min, and subsequently blocked with 1% donkey serum albumin. The cells were then incubated with primary antibodies (1:100 dilution) in a humid chamber overnight at 4 °C. Following three washes, the specimens were incubated with an Alexa Fluor-conjugated secondary antibody (Jackson ImmunoResearch Laboratories Inc) for 1 hour, followed by DAPI (Roche) staining for 30 min at room temperature. Finally, the samples were subjected to confocal microscopy.

### siRNA and lentivirus

siRNA control (siCtrl) and siRNA targeting the gene encoding MSX2 (siMSX2) were purchased from Sigma-Aldrich and transfected into cells using jetPRIME reagents (Polyplus-transfection) following the manufacturer’s instructions. The sequence of siMSX2 is 5’-AGCGCAAGUUCCGUCAGAAAC-3’, while the sequence of sictrl is 5’-UUCUCCGAACGUGUCACGUTT-3’. For lentivirus-delivered RNA interference, negative control (shCtrl) and shRNA targeting MSX2 (shMSX2) were purchased from Shanghai GeneChem Company, with shMSX2 designed to target the same sequence as siMSX2 mentioned above.

For lentiviral production and infection, the empty vector (GV341) was used as a control (OEVector). ectopic expression of MSX2 was used as overexpression MSX2 (OEMSX2), and they induced by lentivirus infection as above were purchased from Shanghai GeneChem Company.

### RNA isolation and quantitative real-time PCR (qPCR)

Total RNA was extracted following the manufacturer’s guidelines using RNA Trizol (TAKARA). A quantity of one microgram of extracted total RNA was converted into cDNA using the PimeScript RT-PCR kit (TAKARA), and qPCR analysis was performed on the LightCycler96 system (Roche) employing the SYBR Premix Ex Tq kit (TAKARA). The levels of amplified mRNA were normalized against 18S mRNA. The primers utilized are detailed in Table S1. Data presented in this study were obtained from a minimum of three independent experiments.

### Chromatin immunoprecipitation (ChIP) and ChIP re-IP

ChIP procedures was conducted as previously described[24].Quantitative PCR (qPCR) assays were conducted to evaluate the precipitated genomic DNA samples.All data related to ChIP were derived from three separate experiments. The primer sequences are provided in Table S2.

For ChIP re-immunoprecipitation (re-IP), cross-linked immunocomplexes were first eluted from the initial ChIP by incubating with 10 mM dithiothreitol (DTT) at 37 °C for 30 min. The eluted products were then diluted 50-fold in re-ChIP buffer and subjected to a second round of ChIP using the specified antibodies.

### Cell growth, colony formation, flow cytometric analysis and Drug resistance experiments

T47D and MCF7 cells infected with either OEVector or OEMSX2 lentivirus and with either shCtrl or shMSX2 lentivirus were seeded into 96-well plates at a density of 3000 cells per well with three replicates for each condition. The cells were treated with estradiol, Tamoxifen (Abmole, Cat# M7353), Fulvestrant (Abmole, Cat# M1966), or Letrozole (Abmole, Cat# M3699) and cultured for 7 days. Cell viability was assessed at various time points using the MTS assay (Promega, Cat# G3580), measuring absorbance at 490 nm.

For the colony formation assay, cells were seeded in 35 mm dishes and cultured in medium supplemented with E2, Tamoxifen, Fulvestrant, or Letrozole at specified concentrations for two weeks. After the incubation period, the cells were fixed with 4% paraformaldehyde for 15 min and subsequently stained with crystal violet to quantify the number of colonies.

For cell cycle analysis, cells were harvested with EDTA-free trypsin two days after treatment with 100 nM E2 and fixed in 75% ethanol at -20 °C overnight. The following day, the cells were washed once with ice-cold PBS, and nuclear DNA was stained with propidium iodide in the dark for 15 minutes. The samples were then analyzed using a flow cytometer (Becton Dickinson).

### Animal experiments and Xenograft tumor studies

All mouse experiments were conducted in strict accordance with the guidelines of the Institutional Animal Care and Use Committee (IACUC) of China Medical University (Ethics Approval Number: CMU20231096). MCF-7 and MCF-7 TMR cells were stably transduced with empty vector and overexpression MSX2 (OEMSX2) lentivirus, respectively, serving as the blank and control groups. Each cell type was suspended in 50 µl of culture medium and mixed with 50 µl of Matrigel (BD Biosciences), then subcutaneously injected into 4-week-old female BALB/c nude mice (Vital River Laboratory) at a concentration of 5 × 10^6^ cells per mouse. The drug-resistant cell group was further divided into control and treatment groups, with the treatment group receiving tamoxifen citrate (10 µg per tablet) every three days. Tumor sizes were recorded by electronic caliper every five days, and tumor volumes were calculated according to the formula: Volume = (width^2^ × length) /2. 30 days after injection, mice were killed by cervical dislocation in keeping with the policy of the humane treatment of tumor bearing animals. After harvesting the xenograft tumors, a part was fixed with formalin for IHC stain.

### Statistical analysis

Statistical analyses were performed using GraphPad Prism 8.02 and SPSS 27.0 software. For all in vitro and in vivo experiments, a two-tailed Student’s t-test was used to calculate P values, with bar graph data presented as mean ± standard deviation (SD) from at least three biological replicates. In animal experiments, mice were randomly assigned in an unbiased manner, and researchers were not blinded during the experiments. The sample size was designed to ensure a 90% probability of detecting statistically significant differences in efficacy between the empty vector and overexpression MSX2 (OEMSX2) groups at a significance level of P < 0.05. All treated experimental units were included in the analysis. The correlation between MSX2 expression and clinical parameters was assessed using the chi-square test. Overall survival curves were generated using the Kaplan-Meier method and compared using the log-rank test.

## Declarations

### Ethics approval and consent to participate

All specimens were collected with informed consent from patients and approved by the Ethics Committee of China Medical University (no. 2025-032). The experiments strictly adhered to the ethical principles outlined in the WMA Declaration of Helsinki and the Belmont Report by the U.S. Department of Health and Human Services.

All mouse experiments were conducted in strict accordance with the guidelines of the Institutional Animal Care and Use Committee (IACUC) of China Medical University (Ethics Approval Number: CMU20231096). All experiments were performed in accordance with the approved protocol and other relevant guidelines and regulations.

### Consent for publication

All authors have agreed with publishing this manuscript.

### Availability of data and materials

All the data can be obtained by contacting the corresponding author.

### Authors’ contributions

**JWF**: Conceptualization, Investigation, Validation. **KZ**: Data curation, Formal analysis. **AQW**: Data curation, Formal analysis. **HMS**: Data curation, Formal analysis. **ZNJ**: Investigation. **ZL**: Investigation. **SF**: Investigation. **XDS**: Investigation. **YB**: Investigation. **MCH**: Investigation. **WSL**: Investigation. **MLW**: Investigation. **DJY**: Investigation. **JW**: Investigation. **WL**: Investigation. **LX**: Writing – review & editing, Funding acquisition. **CYW**: Writing – review & editing, Funding acquisition. **SLW**: Conceptualization, Writing – original draft, Writing – review & editing. **YZ**: Conceptualization, Writing – review & editing, Funding acquisition.

## Acknowledgements

We appreciate Dr. Zhizong Sun (Donggang Stomatological Hospital, Dandong, China), Dr. Yunlong Huo and Dr. Ye Kang (Shengjing Hospital of China Medical University, Shenyang, China) for helpful technique assistance. We thank Dr. Shigeaki Kato (Soma Central Hospital, Fukushima, Japan) for giving us the valuable suggestion and comments for the whole work, and providing a pEREtk-Luc reporter vector, expression plasmids for ERα, and its truncated mutants. We thank Dr. Yujie Sun (Nanjing Medical University, Nanjing, China) for generously providing MCF-7 cells, which were purchased from American Type Culture Collection. We thank Prof. Qinghuan Xiao (the School of Pharmacy, China Medical University, Shenyang, China) for generously providing the tamoxifen-resistant MCF-7 (MCF-7 TMR) cell lines.

